# Computing Linkage Disequilibrium Aware Genome Embeddings using Autoencoders

**DOI:** 10.1101/2023.11.01.565013

**Authors:** Gizem Taş, Timo Westerdijk, Eric Postma, Project MinE ALS GWAS Consortium, Jan H. Veldink, Alexander Schönhuth, Marleen Balvert

## Abstract

**Motivation:** The completion of the human genome has paved the way for genome-wide association studies (GWAS), which have already succeeded in explaining certain proportions of heritability. GWAS are not optimally suited to detect potential non-linear effects in disease risk, possibly hidden in non-additive interactions (epistasis). Alternative methods for epistasis detection using e.g. deep neural networks are currently under active development. Despite their promise to scale to high-dimensional data, deep neural networks (DNNs) are constrained by finite computational resources, which can be rapidly depleted due to increasing complexity with the sheer size of the genome. Besides, the curse of dimensionality complicates the task of capturing meaningful genetic patterns for DNNs; therefore calls for precaution in the form of dimensionality reduction.

**Results:** We propose a method to compress genotyping data, involving single nucleotide polymorphisms (SNPs), while leveraging the LD structure in the genome and preserving non-linear relations among variants. This method involves clustering correlated SNPs into haplotype blocks and training per-block autoencoders that are able to learn a compressed representation of the block’s relevant genetic content. We provide an adjustable autoencoder design to accommodate diverse blocks and to bypass extensive hyperparameter tuning. We applied this method to genotyping data from Project MinE which involves a total of 23,209 ALS cases and 90,249 healthy controls. We compressed the haplotype blocks of an entire chromosome using our autoencoder-based approach, and show that this leads to more than 99% average test reconstruction accuracy - i.e. minimal information loss - while compressing the input to nearly 10% of the original size. We demonstrate that haplotype-block based autoencoders outperform their linear alternative Principal Component Analysis (PCA) by approximately 3% chromosome-wide accuracy of reconstructed variants. To the extent of our knowledge, our approach is the first to simultaneously leverage genome haplotype structure and deep neural networks for dimensionality reduction of genetic data.

**Availability and Implementation:** Data used in this study are available for academic use through the Project MinE Consortium at https://www.projectmine.com/research/data-sharing/. Data accessibility is contingent upon any terms or requirements specified by the source studies. Codes for data compression are available at https://github.com/gizem-tas/haploblock-autoencoders.

## Introduction

In recent years, advances in next-generation sequencing technologies have enabled sequencing of the entire human genome, which has brought us closer to understanding the genetic architecture of complex diseases. Identifying the genetic factors responsible for the variation in disease disposition of individuals holds the promise of developing treatments for currently incurable diseases [Manolio et al., 2009].

High-throughput genome sequencing has paved the way for genome-wide association studies (GWAS) to seek relations between disease status and a large number, sometimes above a million, of genetic variants [Hardy and Singleton, 2009]. Usually, the phenotypic variance explained by genetic variants falls behind its expected portion to be explained, by a difference referred to as the *missing heritability* [Maher, 2008, Manolio et al., 2009]. Taking amyotrophic lateral sclerosis (ALS) as an example, heritability is estimated to be around 45% [Ryan et al., 2019, Trabjerg et al., 2020], which is only partially attributed to the significant loci identified by the latest GWAS [van Rheenen et al., 2021]. In case of many complex diseases, the risk loci from GWAS typically explain a limited amount of the genetic variance; when in fact most heritability may be disguised in moderate or even small effects of allegedly non-significant common loci [Shi et al., 2016, Boyle et al., 2017]. Furthermore, non-additive interactions (i.e. epistasis) between already identified loci might account for part of the missing heritability [Zuk et al., 2012, Blanco-Gómez et al., 2016]. All in all, solving the mysterious genetics of complex diseases seems only likely when joint analysis of many variants, ideally the entire genome, is feasible.

The increasing availability of large-scale raw genotype data has boosted interest in deep learning techniques that are able to scale to high-dimensional data [Wainberg et al., 2018]. However, a deep neural network’s (DNN) performance is still limited by available computational resources, which could easily be exhausted given the exponential increase in the number of network parameters due to the high input dimensionality resulting from the sheer size of genotyping data. In genomics, the number of genetic variants can amount to millions, typically much higher than the number of samples due to practical, technical and cost-related limitations [Gazestani and Lewis, 2019]. As follows from those dimensions, processing large genetic input in its raw state would already be computationally too heavy, even for a shallow neural network with one hidden layer, let alone a deeper one with several layers stacked.

The challenge, widely known as the *curse of dimensionality*, can manifest itself in issues beyond computational complexity [Donoho et al., 2000]. For example, SNP data can partially be sparse due to rare occurrences of some variants in the population. Data sparsity can jeopardize generalization of learning models because of a high proportion of zeros, such that the model’s predictive ability suffers from overfitting [Altman and Krzywinski, 2018]. Alternatively the model could overlook the predictive power of sparse features while giving more weight to denser ones. Altogether, the *curse* of high dimensionality makes finding meaningful patterns harder for DNNs and possibly induces spurious genotype-phenotype associations. Dimensionality reduction may alleviate this problem.

Researchers have suggested remedies for handling sparse input data in DNNs, such as regularizing the input through a lasso penalty [Feng and Simon, 2017]. Alternatively, one could rely on biological priors to address the sparsity in high-dimensional features by eliminating redundant features. The phenomenon of alleles at different loci being associated in a non-random manner is termed *Linkage Disequilibrium (LD)* [Slatkin, 2008]. High or complete LD between SNP pairs is often pointed out as the source of redundancy in large-scale genomic data and removing loci based on their pairwise correlation is a prevalent practice called LD pruning [Calus and Vandenplas, 2018]. By definition, rare variants with low minor allele frequencies are subject to neither complete nor high LD, therefore LD pruning makes practical sense solely for common variants. Besides, such pre-selection methods have the risk of missing some complex higher-order interactions between genetic loci. Reducing the data dimensionality sufficiently to make the models computationally feasible would require strict LD thresholding, which would eventually sweep away possibly disease-causing interactions between genetic loci.

Another use of LD is to segment genome-wide information into non-overlapping substructures by quantifying the correlation between genetic variants. It is not unprecedented to divide the genome into sub-groups of genetic markers prior to dimensionality reduction, as a refuge from multi-collinearity. Hibar et al. have summarized correlated (projected) SNP information within each gene to orthogonal principal components that explain 95% of the total variance per gene [Hibar et al., 2011]. Li et al. address such a collinearity through a local window approach, scanning each chromosome in order to form high-LD clusters of genetic loci [Li et al., 2018]. Information contained in each of these non-overlapping clusters is then summarized into a number of independent principal components, chosen customarily based on a cumulative variance threshold. Despite its usefulness in disposing of the redundancy in sparse and large-scale data, Principal Component Analysis (PCA) is a linear transformation which is only able to preserve the original variance as long as the input features, in this case SNPs, can be assumed to interact linearly [McVean, 2009, Alanis-Lobato et al., 2015]. This would not be the ideal way out of the dimensionality issue when missing heritability is concerned, for bearing the risk of obscuring non-linear relationships between SNPs.

Autoencoders, on the other hand, offer a compelling solution due to their ability to compress high-dimensional genotype data effectively while preserving complex and non-linear patterns in the data [Bank et al., 2020]. First designed as a neural network trained to reconstruct its input in the output layer [Rumelhart et al., 1986], autoencoders were advertised as more powerful yet costly non-linear generalizations of PCA transformation [Hinton and Salakhutdinov, 2006, Fournier and Aloise, 2019]. An autoencoder is also capable of learning an abstract representation of the data internally. These concurrent utilities of autoencoders helped diversify their intended use cases, among which dimensionality reduction has been a prominent one [Goodfellow et al., 2016].

Typically, an autoencoder comprises two components: the *encoder* network, which learns the data relationships and compresses the input into a lower-dimensional bottleneck, and the *decoder* network, which utilizes the abstract representation in the bottleneck to reconstruct the original input data. Learning is achieved through minimizing a loss function, mostly known as the *reconstruction loss*, which aims to penalize the divergence between the input and the output vectors [Goodfellow et al., 2016]. Next, meaningful input patterns captured during training can be extracted from the low-dimensional bottleneck to launch downstream tasks.

Above all, our aim is to compress high-dimensional genotype data in a way that preserves genetic patterns. Considering the uncharted genetic nature of complex diseases such as ALS, we ought to take the quest further into scalability, non-linearity and reproducibility. We strive to find the balance between the curse of dimensionality and unwanted loss of SNP information. Our strategy is to leverage the correlation between SNPs, in other words LD, so as to maximize compression across non-overlapping segments of the genome. We hereby make use of haplotype blocks, i.e. sections of the genome that have high internal LD [Barrett et al., 2005], as a way of drawing the boundaries between clusters of correlated SNPs. Next in our workflow, illustrated in Figure 1, comes the compression of each haplotype block in a low-dimensional space through deep autoencoders. We hypothesize that exploiting the local correlation among SNPs could secure dual benefits, as it would not only mitigate the loss of valuable information but also overcome the obstacles posed by the curse of dimensionality.

**Fig. 1:**
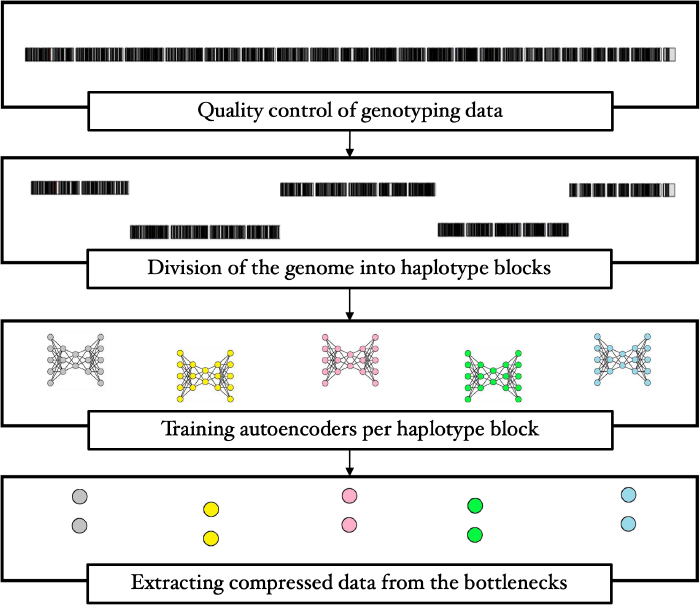
An overview of the workflow. Haplotype block estimation is followed by per-block autoencoder training, where each block is compressed in the corresponding bottleneck layer.

In order to achieve the proposed compression, many haplotype blocks will need to be compressed and many autoencoders will need to be trained in parallel, namely one for each haplotype block. Given the variety of these blocks, optimization of each network’s hyperparameters would consume time and resources beyond practicality, even feasibility. Therefore in this work, we seek a standardized way of building an autoencoder, which manages the trade-off between the rate of input compression and the reconstruction accuracy. We investigate the connection between certain characteristics of haplotype blocks and their corresponding optimal autoencoder configurations. Furthermore, we discuss the opportunities for favouring computational efficiency over marginal returns in performance. Following the versatile analysis, we finalize by compressing an entire chromosome and assess the performance of our framework in terms of both quantified reduction and conservation of significant data patterns.

Our contribution is therefore threefold. First, we propose an efficient dimensionality reduction approach that extends beyond linearity based on LD and autoencoders. Second, we provide a standardized way to determine the autoencoder’s architecture for each individual haplotype block. Third, we show that using autoencoders to compress haplotype blocks can achieve a compression of the data to approximately 10% of the original size with a reconstruction accuracy above 99%.

## Materials and methods

### Data description and pre-processing

In this study, we used genotype data from Project MinE, an international effort to collect clinical and genetic data from ALS patients as well as healthy individuals [Project MinE ALS Sequencing Consortium, 2018]. The data contains a total of 23,209 ALS cases and 90,249 healthy controls from Project MinE; including 56,208 males and 57,250 females. The majority of the participants are of European descent. Diagnosis of ALS patients was carried out according to the revised El-Escorial criteria [Brooks et al., 2000]. Preceding the most extensive GWAS of ALS carried out so far, participants were gathered in cohorts according to their genotyping batch, where they passed individual and variant-level quality control [van Rheenen et al., 2021]. Subsequently, these cohorts were merged as to form five strata based on their genotyping platforms and were once again subjected to quality control and imputation. We then merged four of those strata, which contain 100,049 individuals from European descent. The fifth stratum will be used for testing.

For this study, we only allowed for common SNPs with minor allele frequencies (MAF) above 0.01 and excluded all the SNPs with non-zero missing genotype rates, which could induce bias, using PLINK 1.9 [Marees et al., 2018, Purcell et al., 2007]. As a result of these consecutive quality control and filtering steps, 6,546,842 autosomal SNPs were kept for the analysis. All computations were executed using the University Medical Center Utrecht’s High-Performance Compute (HPC) cluster facilities.

To format the genotype data into data that is usable by autoencoders, we encoded our SNP input to single allele dosage values of 0, 1 or 2, using PLINK 1.9. We resorted to *additive recoding* in particular, which means that allelic dosage values are obtained by counting the minor alleles at a specific locus per person. This conversion generates a raw tabular data format where each SNP is represented by a unique column.

### Parsing the human genome into haplotype blocks

Combinations of alleles or genomic variants, such as polymorphisms, that are inherited together from a single parent constitute a haplotype [National Human Genome Research Institute, 2022]. Constituents of a haplotype reside near each other on a chromosome [International HapMap Consortium, 2005]. The term haplotype block is then used to describe a bounded genomic region which harbours only a few distinct haplotypes [Wall and Pritchard, 2003]. We make use of the procedure suggested by Gabriel et al. for estimating boundaries of haplotype blocks [Gabriel et al., 2002], which is implemented in the widely known Haploview algorithm [Barrett et al., 2005]. The algorithm delimits a haplotype block over a region where only a minor share of the SNP pairs exhibit signs of historical recombination. The authors hypothesize that haplotype block boundaries are considerably aligned across different populations, as are certain haplotypes residing in those blocks. This argument ensures that haplotype block-based dimensionality reduction is not too susceptible to the underlying population structure and can be standardized as the first step of our methodology without a substantial need to account for genetically diverse cohorts.

We estimated haplotype blocks using PLINK 1.9 [Purcell et al., 2007], which implements Taliun et al.’s solution to the Haploview algorithm [Taliun et al., 2014]. In order to obtain lengthy, that is more crowded, blocks - which is advantageous for dimensionality reduction - we calibrated Haploview’s default parameters. Namely, we tailored the confidence interval for strong LD between 0.5 and 0.85, as well as the upper bound for historical recombination, which was set to 0.7 instead of 0.9. Furthermore, we extended the SNP window option from 200 Kb to span over 10 Mb of the genome at once. Such adjustments have indeed helped elongating the haplotype blocks.

### Autoencoders

Next, autoencoders come into play to learn lower-dimensional representations of each haplotype block. One by one, the models are trained in order to reconstruct their highly correlated SNP input, such that valuable genetic patterns can be compressed in the information bottleneck. We built and trained our models using the TensorFlow library [Abadi et al., 2015] in Python 3 [Van Rossum and Drake, 2009].

### Model architecture and configuration

Our approach entails building and training autoencoders individually for each haplotype block. Because of the large number of haplotype blocks, this process needs to be highly standardized. Due to the diversity of the number of SNPs in each block, it would not be ideal to train models with identical architectures for each block. On the other hand, designing neural network architectures for each individual haplotype block would not be feasible. Instead, we propose a flexible mechanism, to piece the autoencoder together based on three hyperparameters, i.e. *shape, number of hidden layers* and *bottleneck dimension*, which determines the achieved rate of compression in the end.

Each autoencoder consists of sequential fully-connected layers, such that the dense connection between consecutive layers can compress the input into a lower dimension at minimum information loss, eliminating redundancy. The number of hidden layers *hl* corresponds to the number of layers between both the input and output layers and the bottleneck, hence the depths of the encoder as well as the decoder. The total number of layers in the autoencoder is 2 *· hl* + 3.

The number of neurons in each layer *l* is determined by the hyperparameter *shape*, and is computed using the bottleneck dimension (*bn*), the number of hidden layers (*hl*), and the slope of the encoder and decoder geometries (*p*). *p* can be 0 or 0.5, corresponding to the shape types *rectangular* or *elliptic*, respectively, see Figure 2. The number of neurons in layer *l* is then calculated by the function *n*(*l*):

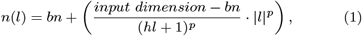

where

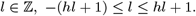

The number of neurons in the input and output layers are given by *n*(*-*(*hl* + 1)) and *n*(*hl* + 1) respectively and are both equal to the input dimension. The bottleneck dimension is given by *n*(0) and is equal to *bn*. The number of neurons in a layer should always be a positive integer, otherwise *n*(*l*) is rounded to the nearest one. The weights of the hidden layers are initialized with He uniform variance scaling initializer [He et al., 2015] and we use the Leaky Rectified Linear Unit activation function (Leaky ReLU) in every layer but the output [Maas, 2013]. The output activations should range from 0 to 2, to approximate our recoded SNP values. Hence, we fashioned the following custom activation function, *r*(*x*), based on the tangent hyperbolic activation function (*tanh*(*x*)), whose output values are bounded between 0 and 2 instead of the original range from -1 to 1:

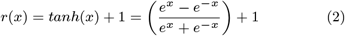

**Fig. 2:**
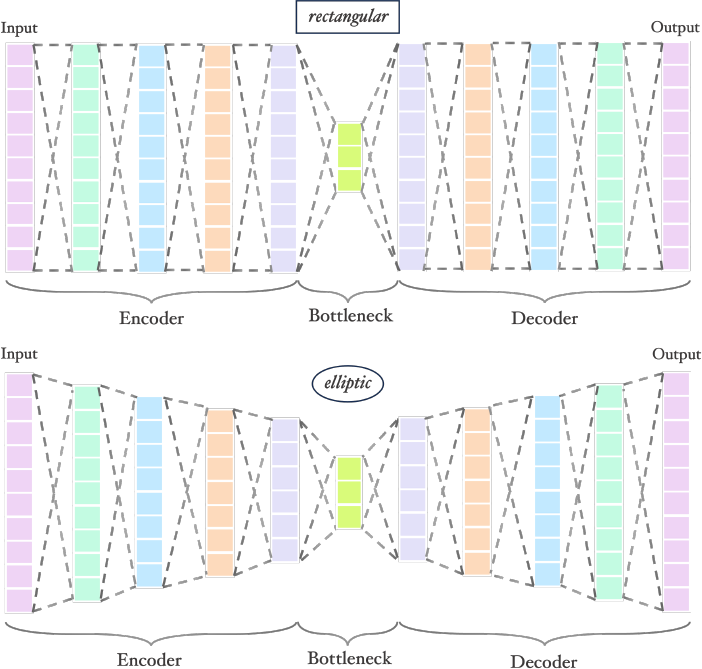
Visualizations of rectangular and elliptic autoencoders. Both networks have the same number of inputs, number of hidden layers and bottleneck size (*inputs* = 10, *hl* = 4, *bn* = 3) and only differ by the slope *p*, see also equation (1).

### Training and evaluation

We preserved 100,049 individuals (20,955 cases and 79,094 controls) in four strata (approximately 88.18% of data) as our training and validation subsets while the remaining stratum of size 13,409 (2,254 cases and 11,155 controls) is set aside for testing. All the hyperparameter tuning experiments were conducted exclusively on the training subset excluding the test stratum.

Since the variants’ allelic dosage values (0, 1, 2) are of ordinal nature, we used Mean Squared Error (MSE) to quantify the reconstruction loss. Autoencoder weights are optimized using the Adam algorithm where the learning rate is initialized as 0.0001 [Kingma and Ba, 2017]. On each haplotype block, autoencoders were trained for 50 epochs with a batch size of 32.

Two metrics are indicative of how well every single autoencoder performs: the *reconstruction loss* and the *SNP reconstruction accuracy*. The former is calculated with the MSE function as a measure of the discrepancy between the input and the output values. The latter assesses the accuracy of individual predictions and is computed by rounding the predicted values of the SNPs to the nearest integer and then comparing those to the input values one by one. Thus, the SNP reconstruction accuracy is the percentage of correctly predicted SNP dosages.

Moreover, we are interested in assessing the extent to which the autoencoders are able to compress the haplotype blocks. By definition, the dimension of the bottleneck layer is equivalent to the reduced data size, and its ratio to the original input size, i.e. the number of SNPs in a haplotype block, yields the compression ratio:

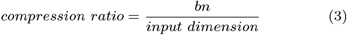

### Hyperparameter tuning

We adopted an adjustable structure defined through 3 hyperparameters (*shape, hl* and *bn*) which ideally need to be optimized for each haplotype block individually. While tuning the hyperparameters of each model seems theoretically possible, it would require an extensive memory and computation time.

In our case, the marginal gain from an extensive hyperparameter tuning procedure may not be as favorable as economizing on computational resources. To bypass the burden of tuning every autoencoder, we first selected a diverse subset of 221 haplotype blocks across 22 chromosomes (autosomes) and optimized hyperparameters for each of these blocks individually. To ensure that we select haplotype blocks of various lengths, in each chromosome, we grouped haplotype blocks in bins based on their sizes and randomly sampled one block per bin. This way, some of the blocks with a high number of member SNPs, which would have been outliers in a distribution-based selection, were represented in our subset.

For each of the selected blocks, we carried out a grid search over a parameter space in which the values for *bn* range from 1 to 10, *hl* from 1 to 5, and the shape of the autoencoders is either *elliptic* or *rectangular*. During this process, we used 5-fold cross-validation with 100,049 training individuals to ensure that the testing stratum remained unseen by any of the trained autoencoders. Meanwhile we recorded both MSE losses and SNP reconstruction accuracies of the 5-fold validation subsets for evaluation.

For downstream analyses, the bottleneck layer of each autoencoder outputs a compressed version of the original haplotype block input. Choosing the number of neurons in the bottleneck layer is a way of predetermining the level of dimensionality reduction. We treat the bottleneck size as a measure for the achieved compression, as well as a hyperparameter to be optimized. Specifically, we ran the grid search for 221 haplotype blocks in our subset, to find the best possible combinations of shape and hidden layers per block, under fixed bottleneck sizes. We repeated the search for every bottleneck size from 1 to 10 to obtain optimal configurations while controlling for the compression level.

## Results

### Descriptive statistics of the haplotype blocks

Due to our method’s dependency on linkage disequilibrium for defining haplotype blocks, we ruled out rare genetic variants from the original data with a minor allele frequency (MAF) cutoff of 0.01. We have clustered the remaining 6,546,842 common variants into 193,122 haplotype blocks over 22 autosomes.

We observe in Supplementary Figure 8 and in Supplementary Table 2 that the distribution of the sizes of the haplotype blocks is positively skewed in each chromosome, such that the majority of blocks contain relatively small numbers of SNPs while fewer blocks appear more populated.

### The trade-off between compression and reconstruction accuracy

For each haplotype block in our subset of 221, the grid search yielded 10 optimal hyperparameter settings corresponding to 10 values of the bottleneck size. To examine the impact of the compressed input dimensionality on the SNP reconstruction accuracy, we averaged the optimal validation accuracies over all the haplotype blocks for each of the 10 different bottleneck sizes. We also estimated the 95% confidence intervals around these optimal validation accuracies, which can provide insight into the reliability of the SNP accuracy estimates across different bottleneck sizes, see Figure 3.

**Fig. 3:**
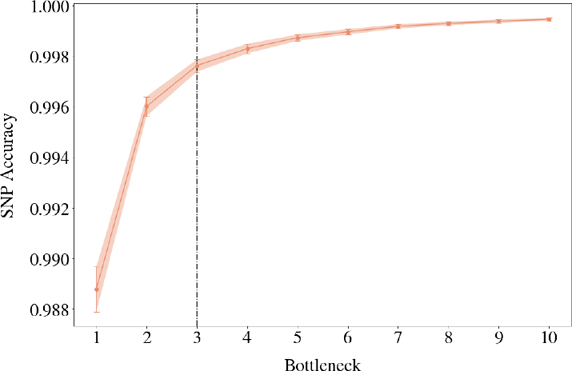
Elbow plot displaying the validation reconstruction accuracies of the best models across bottleneck values from 1 to 10, averaged over 221 haplotype blocks along with 95% confidence intervals. The black line indicates the elbow point observed at 3, beyond which the incremental improvement in accuracy diminishes.

A wider bottleneck can lead to higher SNP reconstruction accuracies by allowing for more detailed representations, as supported by our findings in Figure 3. Also, the confidence intervals become larger on decreasing width of the bottleneck layer. Given the constant sample size of 221, a larger confidence interval necessarily indicates increasing standard deviation, thus a higher degree of uncertainty associated with the average SNP accuracies. Finding the optimal bottleneck dimension involves striking a balance between a high level of compression and maintaining sufficient information for accurate reconstruction. For this, a trade-off point, where the diminishing return in SNP reconstruction accuracy is no longer worth widening the bottleneck, can be found at the bottleneck size of 3, see Figure 3. The average SNP accuracy improves only marginally (by 0.0016) with an additional bottleneck dimension between 2 and 3. Below we explore when we can prioritize compression by pushing the elbow point.

### The influence of haplotype block characteristics on reconstruction accuracy

Every haplotype block can harbour a different number of SNPs. This property can also be implicitly linked to the physical length of the block, hence the extent of genetic variation within as supported by Supplementary Figure 9. Figure 4 illustrates how the relationship between the block size and SNP accuracy is affected under different scales of compression. Evidently, the more rigorous the compression becomes, the less robust the SNP accuracy is to increasing block sizes.

**Fig. 4:**
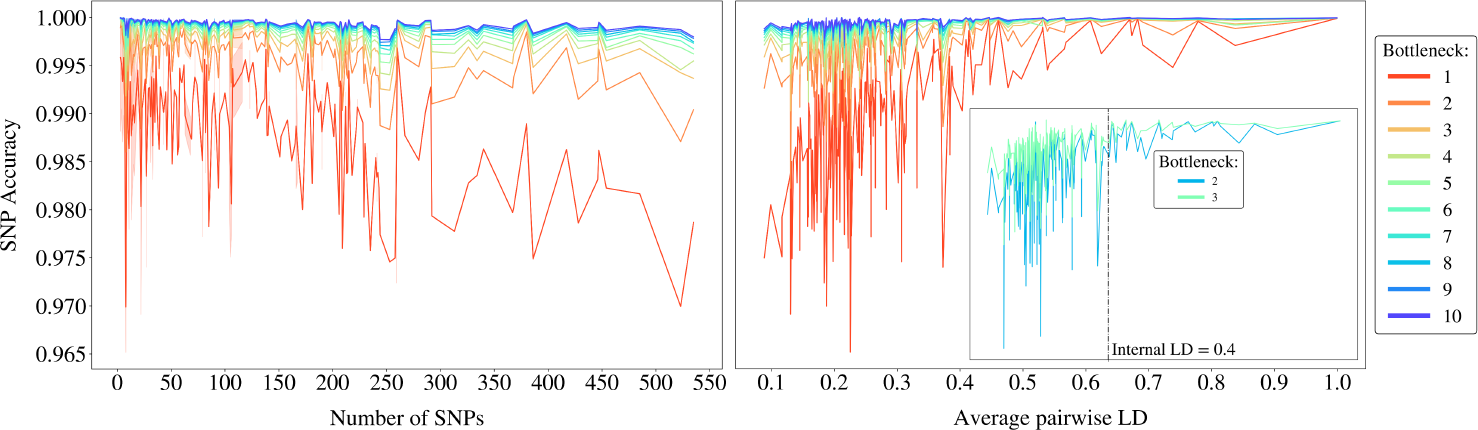
The highest validation accuracies obtained from the grid search for each block, with bottleneck values ranging from 1 to 10. On the left, the accuracies are plotted against the number of SNPs in each block, whereas the x-axis on right side displays the average pairwise LD. The inset zoomed-in plot on the right focuses on bottleneck values 2 and 3, offering a clearer comparison and highlighting the LD threshold at 0.4.

Haplotype blocks can otherwise be assessed by quantifying their internal genetic variation using *the average pairwise linkage disequilibrium (LD)*. The higher the average pairwise LD within a block, the more similar the haplotypes are to each other, indicating lower variation. Overall, a higher degree of compression triggers the sensitivity of SNP accuracy to the variation in blocks, see Figure 4. The positive correlation between the accuracy and average pairwise LD gradually becomes weaker from a bottleneck of size 1 to a bottleneck of size 10.

Given the elbow shown in Figure 3 at 3 bottleneck nodes and its minor advantage over 2, we zoom in to provide a clearer comparison between them, as given in Figure 4. The SNP accuracy still seems less sensitive to changing internal LD for 3 bottleneck dimensions, although we can observe an average pairwise LD threshold at 0.4, beyond which the difference between 2 and 3 bottleneck nodes becomes negligible. Below this threshold, haplotype blocks require more nodes in the bottleneck to ensure sufficient flow of information, as they exhibit higher internal genetic variation. Here, we can seize the opportunity to prioritize compression over accuracy by compressing the haplotype blocks to 2 dimensions, if their internal LD is above 0.4. Otherwise we stand by the original elbow at 3.

### The cost of compression

Among the optimal hyperparameter settings for the 221 haplotype blocks, we counted the occurrences of each combination of *bn* values with *hl* and *shape*. The diagonal pattern with darker shades in Figure 5 indicates a negative correlation between the number of hidden layers and the optimal bottleneck sizes amidst the best performing models.

**Fig. 5:**
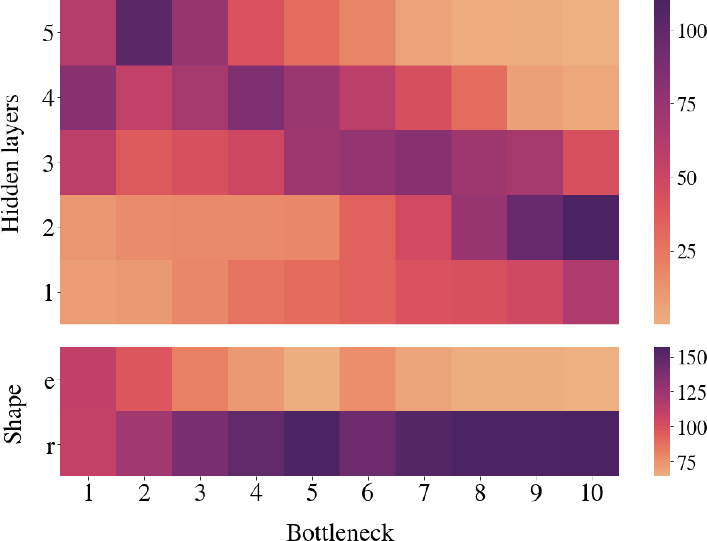
Count of the best models corresponding to the specific pairings of bottleneck sizes with hidden layers on top and model shape on the bottom.

When we track the frequencies of elliptic and rectangular shaped architectures, a large proportion of best models appears to have picked the rectangular shape for nearly all bottleneck sizes, see Figure 5. For the majority of the haplotype blocks, resulting optimal configurations consisted of 5 hidden layers and rectangular shape when the desired bottleneck size is 2 or 3.

To gain further insight into the computational cost incurred by different levels of compression, we monitored the average time (in seconds) taken to train the best performing models across all 5 cross-validation folds during the each block’s grid search. The average of those fitting times across 221 blocks decreases as the bottleneck dimension increases from 2 to 10, see Supplementary Figure 10. Our threefold findings hereby suggest a need for deeper (more hidden layers) and wider (more neurons) autoencoder architectures, hence a proportional rise in computational costs, incurred by higher levels of compression.

### Marginal returns of hyperparameter tuning

To assess the worth of having more hidden layers, we sampled all the haplotype blocks for which the optimal number of hidden layers is 5 and tested for a significant non-zero improvement comparing SNP accuracies resulting from different numbers of hidden layers. Next, we conducted a similar hypothesis test to compare elliptic and rectangular shapes, this time using the blocks for which rectangular is the optimal shape. The results of one-tailed t-tests are given in Supplementary Table 3.

The statistical analysis does not provide sufficient evidence to suggest that using 5 hidden layers instead of 4 hidden layers leads to a significant improvement for bottleneck sizes of 2 and 3, at a significance level of 0.05. Yet in both bottleneck settings, the models achieve significantly higher SNP accuracies when using 4 hidden layers compared to 1, 2, or 3. Furthermore, similarly testing for the difference between rectangular and elliptic shapes resulted in *p*-values above the significance level. The return on the extra computational cost of 5 hidden layers or rectangular autoencoders is thus not evident. Therefore, we conclude that a standard autoencoder featuring elliptic shape and 4 hidden layers can adequately compress diverse haplotype blocks to 2 or 3 dimensions, conditioned on the block’s average pairwise LD.

### Compression of Chromosome 22

Using the chosen autoencoder architecture per haplotype block, we compressed Chromosome 22 - which originally contained 70,247 SNPs clustered in 2,854 blocks - to 7,341 dimensions in total. All autoencoders were trained and evaluated using the same predefined training and test samples. To assess the overall performance, we computed the average losses and accuracies along with their standard deviations across 2,854 autoencoders, see Supplementary Table 4.

In the end, we achieved a dimensionality reduction down to 10.45% of the original input size with an average SNP accuracy of 99.55% on the training samples and 99.56% on the test stratum. The mean MSE losses obtained on the training and test samples were 5.69 × 10^*-*3^ and 4.75 × 10^*-*3^, respectively.

Given the inherent sparsity of genetic input, in the sense that the most common allelic dosage value in the data is 0, the autoencoder may simply output only 0s and still achieve a fair reconstruction accuracy in highly sparse blocks. Therefore, we also evaluate the SNP accuracies of subsets in the data depicted in Figure 6. The reconstruction of 0s yielded the highest average accuracy, followed by 2. The widest spread is obtained for a reconstruction accuracy of 1s.

**Fig. 6:**
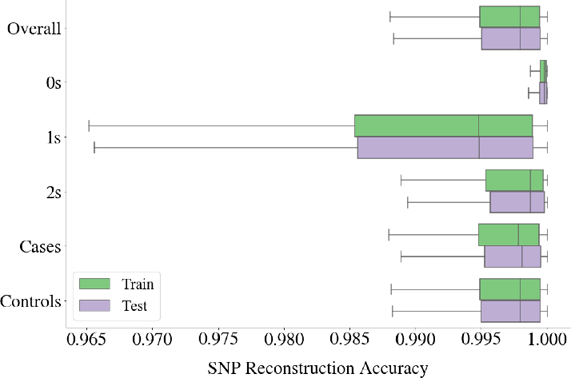
Box plot of SNP reconstruction accuracies obtained from 2854 autoencoders utilized for compression of all the haplotype blocks in Chromosome 22. The accuracies were also calculated separately for 3 categories of genotype dosage values in the data (0s, 1s and 2s), and for 2 phenotypic groups (ALS patients and healthy controls) as displayed on the *y*-axis. The range of SNP accuracies for reconstruction of 1s in the data is the widest, followed by 2s and 0s, with a significantly narrower spread. ALS cases and controls exhibit rather similar accuracy distributions across 2,854 haplotype blocks.

Furthermore, we inspect the SNP accuracies of ALS cases and controls separately to assess whether the disease phenotype impacts the reconstruction performance. The SNP accuracies obtained for both phenotype groups spread within highly similar ranges as can be seen from Figure 6, and their means over all the haplotype blocks differ only marginally, with all values above 99.50% for both training and testing samples, see Supplementary Table 4 for details.

### Comparison with PCA Reconstruction

It is possible to reconstruct the original data from the leading principal components with minimal total squared loss [Plaut, 2018]. For each haplotype block in Chromosome 22, we applied linear transformations using PCA with the same latent space dimensionality as the autoencoder’s bottleneck layer. Here, the orthogonal basis is obtained through the same training data which is used to train the autoencoders, and the test stratum is projected to this basis only for evaluation. Subsequently, we reconstructed the original haplotype blocks from these principal components and calculated MSE losses and SNP reconstruction accuracies by comparing the reconstructed outputs to the inputs, see Supplementary Table 5. Additionally, we calculated the *chromosome reconstruction accuracy* considering the total number of variants correctly predicted in the reconstruction across the entire Chromosome 22 as below:

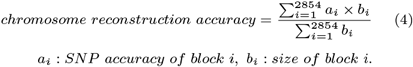

*ai* : *SNP accuracy of block i, bi* : *size of block i*.

Autoencoders have effectively reconstructed 99.63% of the Chromosome 22 variants on the unseen data, while PCA has achieved 96.80%, see Table 1.

**Table 1.** Comparison of PCA and Autoencoder Reconstruction Through Complete Compression of Chromosome 22.

| Method | MSE Loss |  | Chr. Rec. Accuracy |  |
| --- | --- | --- | --- | --- |
|  | Train | Test | Train | Test |
| Autoencoder | $5.69 \times 10^{-3}$ | $4.75 \times 10^{-3}$ | **99.65%** | **99.63%** |
| PCA | $1.54 \times 10^{-2}$ | $1.48 \times 10^{-2}$ | 96.72% | 96.80% |

To pinpoint where this difference originates in, we break the SNP accuracies down based on the allelic dosages (0, 1 or 2) and plot these against block sizes for both methods in Figure 7.

**Fig. 7:**
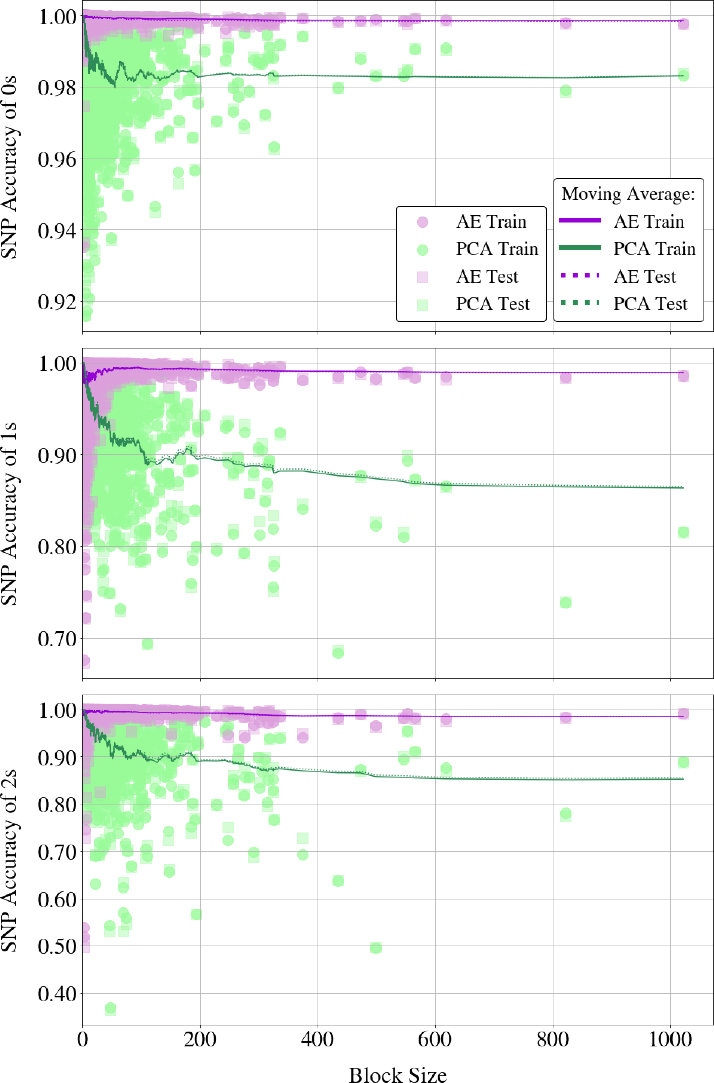
Scatter plots of SNP reconstruction accuracies obtained by both AE and PCA for allelic dosage values of 0, 1, and 2, from top to bottom. The *x*-axes represent block sizes, shared across all plots. Solid lines illustrate the moving average of SNP accuracies for 2 methods on training samples using a window size of 50, while dotted lines depict the same for test accuracies.

Autoencoders consistently outperform PCA across increasing block sizes, maintaining stable performance while PCA’s accuracy declines. As given in Supplementary Figure 11, PCA closely matches autoencoders only for the reconstruction of 1s and 2s in the smallest blocks with less than 10 SNPs, but the disparity in their reconstruction abilities widens evidently with increasing block sizes, see Figure 7.

Furthermore, Figure 7 reveals that the range of the performance gap between 2 methods extends as dosage values become less abundant in the data. For 0s, PCA accuracies remain above 98% and closer to autoencoders, with an erratic performance for smaller blocks. For 1s and 2s on the other hand, PCA accuracies diverge from the autoencoders at a much faster rate, leading to a performance gap that eventually exceeds 10%. Autoencoders excel in accurately reconstructing these scarcer dosage values, also showing robustness to varying block sizes.

Given the sparse aspect of the genetic data, a good enough reconstruction of 0s is necessary but not sufficient to claim that the employed method indeed captured the genuine SNP interactions. This highlights the advantage of autoencoders in capturing genuine SNP interactions in sparse genetic data.

## Discussion

This work introduced a non-linear dimensionality reduction approach to effectively compress massive genotyping data while optimizing the trade-off between retaining meaningful genetic information and low dimensionality. First, we segmented the genome into non-overlapping haplotype blocks to capture the LD structure of the genome. Then, we trained deep autoencoders to compress these haplotype blocks into low-dimensional representations. When these representations are pieced together into a compressed representation of the full genome, the data becomes easier for AI approaches to digest, for example in disease prediction tasks, while allowing for explainability at the haplotype block level. The prominent advantages of our approach can be attributed to several methodological components: (1) leveraging the pairwise correlation between SNPs to facilitate compression using the haplotype block arrangement, (2) retaining the complex interactions between SNPs within those blocks through the layers of the autoencoder, (3) bypassing the burdensome phase of hyperparameter tuning by means of a standardized autoencoder configuration approach and (4) yielding high SNP reconstruction accuracies, indicating minimal loss of SNP information, while compressing large scale genotyping data.

From a population genetics perspective, one would prefer to work with population-specific haplotype block boundaries and training autoencoders specifically tailored to each population since these boundaries are not universally consistent [Zhao et al., 2003, Wall and Pritchard, 2003]. Nonetheless, in cases where certain subpopulations are abundant in the data, autoencoders may be prone to overfitting to dominant genetic patterns driven by those, which would compromise the model’s ability to capture the genetic diversity of underrepresented populations. Instead, we segmented the genome using fixed boundaries to enhance our statistical power and generalization ability.

Imposing a fixed autoencoder architecture to each haplotype block to short-circuit hyperparameter optimization would ignore the diversity of the blocks. In addition to a flexible network design with an adjustable input layer and complexity based on only 3 hyperparameters, our results put forward a decision mechanism for choosing the best configuration applicable to the entire genome. Although this generalized best setting might be sub-optimal for individual blocks, it managed to find the ideal balance between computational efficiency and performance. Our *compromise to optimize* strategy achieved a successful generalization of the autoencoders to an unseen stratum yielding an average test SNP accuracy of 99.63% across 2,854 blocks throughout the entire Chromosome 22.

The dimensionality of each compressed haplotype block can be controlled through the width of the corresponding autoencoder’s bottleneck. There is a trade-off between this width and the SNP reconstruction accuracy, in other words the extent of the information retained in the low-dimensional representations of the blocks. Our methodology allows for customizing the compression settings depending on the scope of downstream tasks. For example, when predicting monogenic or maybe even polygenic traits, assuming that the causal genomic regions can be selected prior to analysis, a wider bottleneck can be chosen on account of a higher accuracy. However, for complex traits with much more intricate genetic architectures according to the hypothesis of the omnigenic model [Boyle et al., 2017], preferably the entire genome should be involved in the analysis. Hence, compressing more genetic information into fewer dimensions becomes the better remedy. As revealed in our results, the reconstruction performance is highly sensitive to the genetic variation (internal LD) covered by the haplotype block notably for lower bottleneck dimensions (below 4). This could mean that a wider bottleneck more steadily ensures the necessary network complexity to reconstruct the genetic input. To provide the desired dimensionality as well as to avoid overcomplicating the networks, we considered LD-thresholding to decide between 2 and 3 bottleneck nodes.

We deduced from the grid search that the lower the dimension of the bottleneck was, the higher the complexity of the best performing models became, hence the run times of these models increased. In principle, grid search appoints the optimal setting only considering the reconstruction performance, and favors a deeper and wider network for the sake of a minor accuracy improvement, regardless of the increased training cost. The accumulated gain from economizing on run time and model complexity during each haplotype block’s compression can lead to enhanced computational efficiency for the entire genome. After statistically assessing the choices made by the grid search, we seized any opportunity to simplify the networks as long as there was no statistical evidence of inferior performance.

Our findings demonstrate the performance advantage of autoencoders over PCA, which seems particularly evident in the reconstruction of 1s and 2s by at least 2% in mean train and test accuracies, see Supplementary Tables 4 and 5. The significance of this statistical advantage can be better grounded with reference to genotyping errors. Even when occurring at a rate below 1%, genotyping errors are deemed non-negligible as they can have serious repercussions for subsequent analyses [Pompanon et al., 2005, Wang, 2018]. Unlike 0, allelic dosage values of 1 and 2 indicate the presence of a minor allele at a particular genetic locus, hence existence of a SNP. As an orthogonal linear transformation, PCA’s limited ability to capture these values hereby hints at the intricacy and non-linearity of the SNP interactions within a block. Despite its low maintenance and cost-effectiveness, breaking the non-linearity in the process and the associated risk of losing epistasis renders PCA unappealing for compressing the genome, an issue that can be overcome with autoencoders.

Although our approach is promising for developing AI-applicable representations of haplotype blocks, there are potential areas for improvement that future work may need to address. Firstly, the foundation of our method is to segment the genome using the pairwise LD between SNPs, which does not directly apply to the rare variants [Zhang et al., 2002]. Besides, the latent space learnt by the autoencoders might not authentically represent the rare variation in the genome due to the low frequency of such variants in the population. Regarding ALS in particular, discovering the pathogenicity of rare variants could play a major role in resolving the genetic mystery of the disease [van Rheenen et al., 2021]. This renders the future development of compression or feature learning strategies focused on rare genetic variants inevitable. Also, estimating the boundaries of haplotype blocks involves presetting a scanning window for SNPs (10 Mb in this study). Therefore, the long-range dependencies between physically distant variants along the genome are not necessarily covered by the haplotype blocks and they might not be captured throughout compression. Fortunately, the corpus for modeling the long-range dependencies in sequential data is progressing fast and showing encouraging outcomes in genomics [Ji et al., 2021, Nguyen et al., 2023] and our compressed representations form suitable inputs for modeling such dependencies at the haplotype block level.

## Conclusions and Future Work

In this article, we presented a dimensionality reduction approach that makes massive genome data compatible with AI techniques. To retain local patterns in the genome, we first partitioned it into non-overlapping clusters of correlated SNPs, forming haplotype blocks. Then, we trained deep autoencoders capable of capturing intricate relationships to compress each haplotype block. Given the unique characteristics of these blocks, we devised adjustable autoencoder architectures using only 3 hyperparameters. Our findings revealed a decision scheme for the optimal hyperparameters, which helped mitigate their resource-intensive tuning process. Evaluation of our method on an entire chromosome showcased the unprecedented potential of leveraging the LD structure of the genome in conjunction with deep autoencoders for data compression, with notably high reconstruction performances. Such a synergy between haplotype blocks and autoencoders holds promise for effectively preprocessing the genome, eventually to demystify the genetic architectures of complex diseases.

Our compression approach enables various possibilities for applications, including an alternative to single SNP-association tests. The compressed representations at the bottleneck can be utilized for conducting association studies at the level of haplotype blocks. While the former might offer higher resolution at the SNP level, the latter can provide additional insights into the genetic architecture of complex traits by capturing the effects of multiple closely linked SNPs [Hirschhorn and Daly, 2005, Hayes, 2013]. Furthermore, drawing inspiration from the denoising applications of autoencoders [Vincent et al., 2010], which reconstruct noiseless targets from perturbed inputs, we believe that our method holds significant potential for performing reference-free imputation of missing SNP values. It is also possible to modify the self-supervised nature of autoencoder training by incorporating a joint loss function, thereby conditioning the learned representations through a supervised task [Dincer et al., 2020]. As part of our plan, we aim to expand our approach to include a complementary task involving ALS classification using the Project MinE dataset, in pursuit of uncovering the missing heritability of the disease.

## Acknowledgments

The authors wish to acknowledge the The High Performance Compute (HPC) facility, as part of the Utrecht Bioinformatics Center (UBC), for providing necessary computing resources for this study. The authors appreciate the support from Kristel van Eijk and Maarten Koyman in ensuring the efficiency of data pre-processing and experimental setup.

## Author contributions statement

G.T. and T.W. conceived and conducted the experiments. M.B, A.S. and E.P. contributed to the study design. J.H.V provided data. G.T. analysed the results and wrote the manuscript. M.B, A.S., E.P. and J.H.V. critically reviewed the analyses and the manuscript.

## References

M. Abadi, A. Agarwal, P. Barham, E. Brevdo, Z. Chen, C. Citro, G. S. Corrado, A. Davis, J. Dean, M. Devin, S. Ghemawat, I. Goodfellow, A. Harp, G. Irving, M. Isard, Y. Jia, R. Jozefowicz, L. Kaiser, M. Kudlur, J. Levenberg, D. Mané, R. Monga, S. Moore, D. Murray, C. Olah, M. Schuster, J. Shlens, B. Steiner, I. Sutskever, K. Talwar, P. Tucker, V. Vanhoucke, V. Vasudevan, F. Viégas, O. Vinyals, P. Warden, M. Wattenberg, M. Wicke, Y. Yu, and X. Zheng. TensorFlow: Large-scale machine learning on heterogeneous systems, 2015. URL https://www.tensorflow.org/. xSoftware available from tensorflow.org.

G. Alanis-Lobato, C. V. Cannistraci, A. Eriksson, A. Manica, and T. Ravasi. Highlighting nonlinear patterns in population genetics datasets. Sci. Rep., 5(1):8140, Jan. 2015.

N. Altman and M. Krzywinski. The curse(s) of dimensionality. Nat. Methods, 15(6):399–400, June 2018.

D. Bank, N. Koenigstein, and R. Giryes. Autoencoders. 2020.

J. C. Barrett, B. Fry, J. Maller, and M. J. Daly. Haploview: analysis and visualization of LD and haplotype maps. Bioinformatics, 21(2):263–265, Jan. 2005.

A. Blanco-Gómez, S. Castillo-Lluva, M. Del Mar Saéz-Freire, L. Hontecillas-Prieto, J. H. Mao, A. Castellanos-Martín, and J. Pérez-Losada. Missing heritability of complex diseases: Enlightenment by genetic variants from intermediate phenotypes. Bioessays, 38(7):664–673, July 2016.

E. A. Boyle, Y. I. Li, and J. K. Pritchard. An expanded view of complex traits: From polygenic to omnigenic. Cell, 169(7): 1177–1186, June 2017.

B. R. Brooks, R. G. Miller, M. Swash, T. L. Munsat, and World Federation of Neurology Research Group on Motor Neuron Diseases. El escorial revisited: revised criteria for the diagnosis of amyotrophic lateral sclerosis. Amyotroph. Lateral Scler. Other Motor Neuron Disord., 1(5):293–299, Dec. 2000.

M. Calus and J. Vandenplas. Snprune: An efficient algorithm to prune large snp array and sequence datasets based on high linkage disequilibrium. Genetics Selection Evolution, 50:34, 06 2018. doi: 10.1186/s12711-018-0404-z.

A. B. Dincer, J. D. Janizek, and S.-I. Lee. Adversarial deconfounding autoencoder for learning robust gene expression embeddings. Bioinformatics, 36(Suppl 2):i573–i582, Dec. 2020.

D. L. Donoho et al. High-dimensional data analysis: The curses and blessings of dimensionality. AMS math challenges lecture, 1(2000):32, 2000.

J. Feng and N. Simon. Sparse-input neural networks for high-dimensional nonparametric regression and classification. arXiv preprint arXiv:1711.07592, 2017.

Q. Fournier and D. Aloise. Empirical comparison between autoencoders and traditional dimensionality reduction methods. In 2019 IEEE Second International Conference on Artificial Intelligence and Knowledge Engineering (AIKE). IEEE, June 2019.

S. B. Gabriel, S. F. Schaffner, H. Nguyen, J. M. Moore, J. Roy, B. Blumenstiel, J. Higgins, M. DeFelice, A. Lochner, M. Faggart, S. N. Liu-Cordero, C. Rotimi, A. Adeyemo, R. Cooper, R. Ward, E. S. Lander, M. J. Daly, and D. Altshuler. The structure of haplotype blocks in the human genome. Science, 296(5576):2225–2229, 2002. doi: 10.1126/science.1069424. URL https://www.science.org/doi/abs/10.1126/science.1069424.

V. H. Gazestani and N. E. Lewis. From genotype to phenotype: Augmenting deep learning with networks and systems biology. Curr. Opin. Syst. Biol., 15:68–73, June 2019.

I. Goodfellow, Y. Bengio, and A. Courville. Deep Learning. MIT Press, 2016. http://www.deeplearningbook.org.

J. Hardy and A. Singleton. Genomewide association studies and human disease. N. Engl. J. Med., 360(17):1759–1768, Apr. 2009.

B. Hayes. Overview of statistical methods for genome-wide association studies (GWAS). In Methods in Molecular Biology, Methods in molecular biology (Clifton, N.J.), pages 149–169. Humana Press, Totowa, NJ, 2013.

K. He, X. Zhang, S. Ren, and J. Sun. Delving deep into rectifiers: Surpassing human-level performance on imagenet classification. In Proceedings of the IEEE international conference on computer vision, pages 1026–1034, 2015.

D. P. Hibar, J. L. Stein, O. Kohannim, N. Jahanshad, A. J. Saykin, L. Shen, S. Kim, N. Pankratz, T. Foroud, M. J. Huentelman, S. G. Potkin, C. R. Jack, M. W. Weiner, A. W. Toga, and P. M. Thompson. Voxelwise gene-wide association study (vgenewas): Multivariate gene-based association testing in 731 elderly subjects. NeuroImage, 56(4):1875–1891, 2011. ISSN 1053-8119. doi: 10.1016/j.neuroimage.2011.03.077. URL https://www.sciencedirect.com/science/article/pii/S1053811911003715.

G. E. Hinton and R. R. Salakhutdinov. Reducing the dimensionality of data with neural networks. Science, 313(5786): 504–507, July 2006.

J. N. Hirschhorn and M. J. Daly. Genome-wide association studies for common diseases and complex traits. Nat. Rev. Genet., 6 (2):95–108, Feb. 2005.

International HapMap Consortium. A haplotype map of the human genome. Nature, 437(7063):1299–1320, Oct. 2005.

Y. Ji, Z. Zhou, H. Liu, and R. V. Davuluri. DNABERT: pre-trained bidirectional encoder representations from transformers model for DNA-language in genome. Bioinformatics, 37(15): 2112–2120, Aug. 2021.

D. P. Kingma and J. Ba. Adam: A method for stochastic optimization, 2017.

Z. Li, P. Kemppainen, P. Rastas, and J. Merilä. Linkage disequilibrium clustering-based approach for association mapping with tightly linked genome-wide data. Molecular Ecology Resources, 18, 04 2018. doi: 10.1111/1755-0998.12893.

L. Maas. Rectifier nonlinearities improve neural network acoustic models. 2013.

Maher. Personal genomes: The case of the missing heritability. Nature, 456(7218):18–21, Nov. 2008.

T. A. Manolio, F. S. Collins, N. J. Cox, D. B. Goldstein, L. A. Hindorff, D. J. Hunter, M. I. McCarthy, E. M. Ramos, L. R. Cardon, A. Chakravarti, J. H. Cho, A. E. Guttmacher, A. Kong, L. Kruglyak, E. Mardis, C. N. Rotimi, M. Slatkin, D. Valle, A. S. Whittemore, M. Boehnke, A. G. Clark, E. E. Eichler, G. Gibson, J. L. Haines, T. F. C. Mackay, S. A. McCarroll, and P. M. Visscher. Finding the missing heritability of complex diseases. Nature, 461(7265):747–753, Oct. 2009.

A. T. Marees, H. de Kluiver, S. Stringer, F. Vorspan, E. Curis, C. Marie-Claire, and E. M. Derks. A tutorial on conducting genome-wide association studies: Quality control and statistical analysis. Int. J. Methods Psychiatr. Res., 27(2):e1608. June 2018.

G. McVean. A genealogical interpretation of principal components analysis. PLoS Genet., 5(10):e1000686, Oct. 2009.

National Human Genome Research Institute. Haplotype. https://www.genome.gov/genetics-glossary/haplotype, 2022. Accessed: 2022-07-21.

E. Nguyen, M. Poli, M. Faizi, A. Thomas, C. Birch-Sykes, M. Wornow, A. Patel, C. Rabideau, S. Massaroli, Y. Bengio, S. Ermon, S. A. Baccus, and C. Ré. HyenaDNA: Long-range genomic sequence modeling at single nucleotide resolution. June 2023.

E. Plaut. From principal subspaces to principal components with linear autoencoders. Apr. 2018.

F. Pompanon, A. Bonin, E. Bellemain, and P. Taberlet. Genotyping errors: causes, consequences and solutions. Nat. Rev. Genet., 6(11):847–859, Nov. 2005.

Project MinE ALS Sequencing Consortium. Project MinE: study design and pilot analyses of a large-scale whole-genome sequencing study in amyotrophic lateral sclerosis. Eur. J. Hum. Genet., 26(10):1537–1546, Oct. 2018.

S. Purcell, B. Neale, K. Todd-Brown, L. Thomas, M. A. R. Ferreira, D. Bender, J. Maller, P. Sklar, P. I. W. de Bakker, M. J. Daly, and P. C. Sham. PLINK: a tool set for whole-genome association and population-based linkage analyses. Am. J. Hum. Genet., 81(3):559–575, Sept. 2007.

D. E. Rumelhart, G. E. Hinton, and R. J. Williams. Learning representations by back-propagating errors. Nature, 323(6088): 533–536, Oct. 1986.

M. Ryan, M. Heverin, R. L. McLaughlin, and O. Hardiman. Lifetime risk and heritability of amyotrophic lateral sclerosis. JAMA Neurol., 76(11):1367–1374, Nov. 2019.

H. Shi, G. Kichaev, and B. Pasaniuc. Contrasting the genetic architecture of 30 complex traits from summary association data. Am. J. Hum. Genet., 99(1):139–153, July 2016.

M. Slatkin. Linkage disequilibrium–understanding the evolutionary past and mapping the medical future. Nat. Rev. Genet., 9(6):477–485, June 2008.

D. Taliun, J. Gamper, and C. Pattaro. Efficient haplotype block recognition of very long and dense genetic sequences. BMC Bioinformatics, 15(1):10, Jan. 2014.

B. B. Trabjerg, F. C. Garton, W. van Rheenen, F. Fang, R. D. Henderson, P. B. Mortensen, E. Agerbo, and N. R. Wray. ALS in danish registries: Heritability and links to psychiatric and cardiovascular disorders. Neurol. Genet., 6(2):e398, Apr. 2020.

W. van Rheenen, R. A. A. van der Spek, M. Bakker, and et al. Common and rare variant association analyses in amyotrophic lateral sclerosis identify 15 risk loci with distinct genetic architectures and neuron-specific biology. Nature Genetics, 53 (12):1636–1648, Dec. 2021.

G. Van Rossum and F. L. Drake. Python 3 Reference Manual. CreateSpace, Scotts Valley, CA, 2009. ISBN 1441412697.

P. Vincent, H. Larochelle, I. Lajoie, Y. Bengio, P.-A. Manzagol, and L. Bottou. Stacked denoising autoencoders: Learning useful representations in a deep network with a local denoising criterion. Journal of machine learning research, 11(12), 2010.

M. Wainberg, D. Merico, A. Delong, and B. J. Frey. Deep learning in biomedicine. Nat. Biotechnol., 36(9):829–838, Oct. 2018.

J. D. Wall and J. K. Pritchard. Haplotype blocks and linkage disequilibrium in the human genome. Nat. Rev. Genet., 4(8): 587–597, Aug. 2003.

J. Wang. Estimating genotyping errors from genotype and reconstructed pedigree data. Methods Ecol. Evol., 9(1):109–120, Jan. 2018.

K. Zhang, P. Calabrese, M. Nordborg, and F. Sun. Haplotype block structure and its applications to association studies: power and study designs. Am. J. Hum. Genet., 71(6):1386–1394, Dec. 2002.

H. Zhao, R. Pfeiffer, and M. H. Gail. Haplotype analysis in population genetics and association studies. Pharmacogenomics, 4(2):171–178, Mar. 2003.

O. Zuk, E. Hechter, S. R. Sunyaev, and E. S. Lander. The mystery of missing heritability: Genetic interactions create phantom heritability. Proc. Natl. Acad. Sci. U. S. A., 109(4):1193–1198, Jan. 2012.

