## Supplementary Data for "Computing Linkage Disequilibrium Aware Genome Embeddings using Autoencoders"

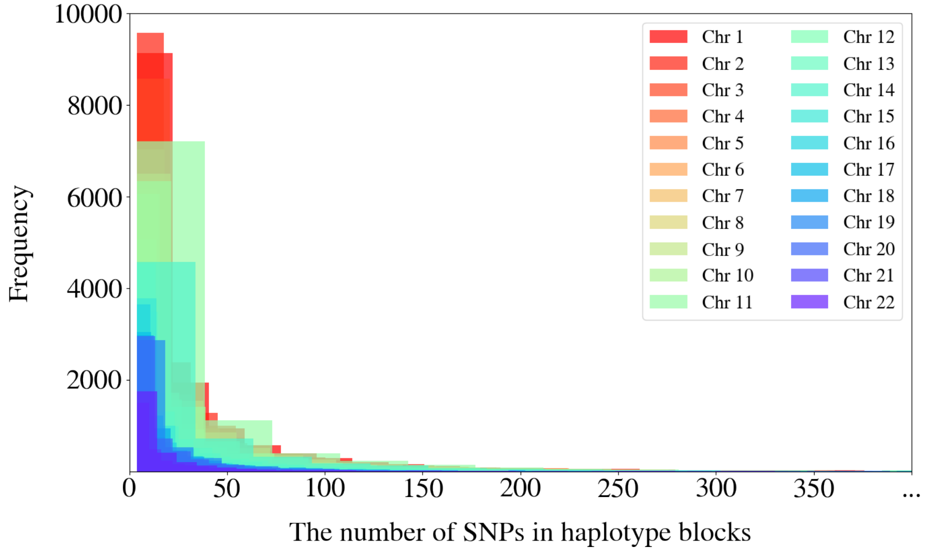


Fig. 8: The distribution of the number of SNPs in haplotype blocks per chromosome. In each chromosome, the sizes of the haplotype blocks have a right-skewed distribution meaning that most blocks contain relatively few SNPs.

**Table 2.** Detailed statistics of haplotype blocks obtained per chromosome. Resulting haplotype blocks differ in size, i.e. the number of SNPs they contain.

| | Total Number of | | Block Size Statistics | | |
|---|---|---|---|---|---|
| Chromosome | Haplotype Blocks | SNPs | Mean | Median | Standard Deviation |
| 1 | 14252 | 507103 | 35.88 | 13 | 66.38 |
| 2 | 16365 | 571572 | 35.19 | 13 | 66.49 |
| 3 | 13778 | 486999 | 35.62 | 13 | 69.28 |
| 4 | 12341 | 486982 | 39.77 | 14 | 72.66 |
| 5 | 11933 | 443075 | 37.44 | 13 | 66.78 |
| 6 | 11080 | 462170 | 42.01 | 15 | 79.43 |
| 7 | 11274 | 381486 | 34.11 | 13 | 61.97 |
| 8 | 11016 | 382486 | 34.99 | 13 | 66.54 |
| 9 | 8961 | 279698 | 31.41 | 12 | 59.27 |
| 10 | 9976 | 343148 | 34.68 | 13 | 64.43 |
| 11 | 9362 | 328257 | 35.35 | 13 | 80.41 |
| 12 | 9006 | 321068 | 35.94 | 13 | 65.64 |
| 13 | 6770 | 251751 | 37.48 | 14 | 69.24 |
| 14 | 6155 | 216676 | 35.46 | 13 | 73.35 |
| 15 | 6543 | 179377 | 27.63 | 11 | 55.37 |
| 16 | 7336 | 177592 | 24.43 | 11 | 42.66 |
| 17 | 5637 | 144668 | 25.89 | 10 | 47.41 |
| 18 | 5903 | 184447 | 31.51 | 12 | 56.06 |
| 19 | 4201 | 103423 | 24.84 | 10 | 48.87 |
| 20 | 5236 | 139515 | 26.87 | 11 | 47.07 |
| 21 | 3143 | 85102 | 27.31 | 11 | 46.74 |
| 22 | 2854 | 70247 | 24.81 | 10 | 50.95 |

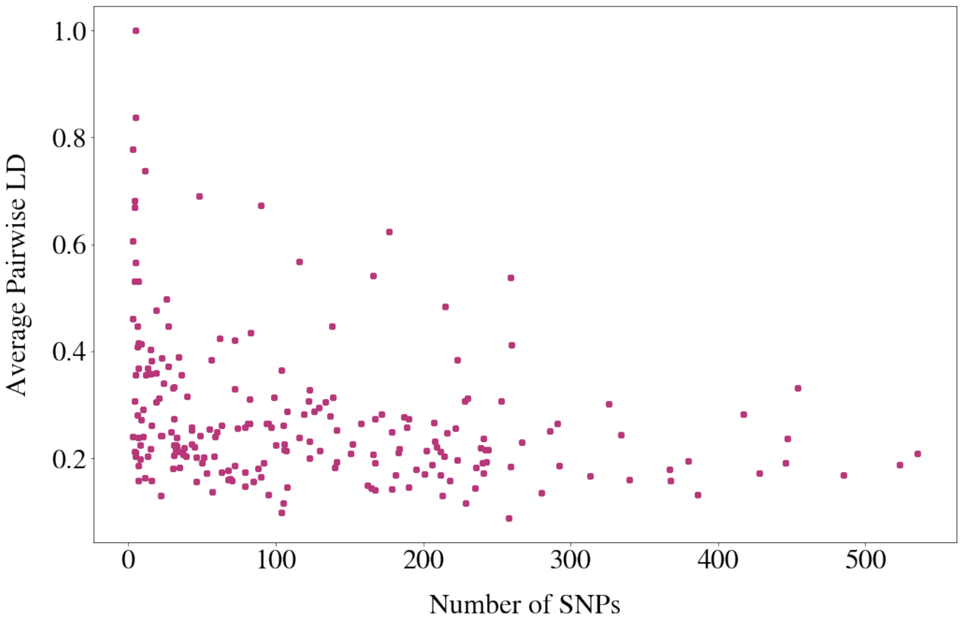


Fig. 9: The weak negative correlation between the average pairwise LD measuring the interval variation and the size of the haplotype blocks quantified by the number of SNPs that they contain. The computed Pearson correlation coefficient of -0.3017 suggests a moderate negative correlation between these two characteristics under consideration.

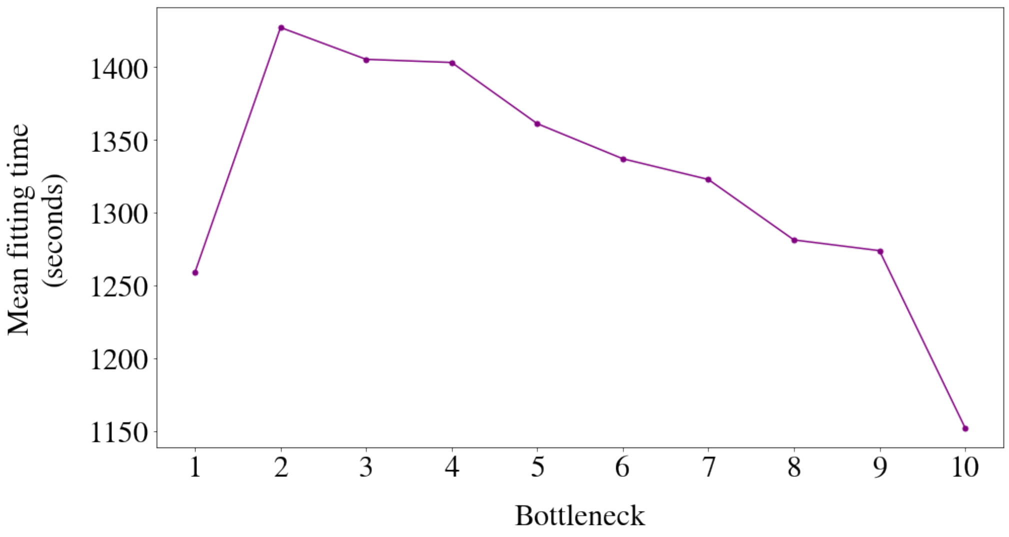


Fig. 10: The mean fitting times (in seconds) obtained by tracking the training times of the best performing models (one for each bottleneck value) across 5 cross-validation folds and computing their average across 221 haplotype blocks. These mean fitting times are plotted against 10 values of the bottleneck. A decreasing trend is observed as bottleneck values progress from 2 to 10, suggesting an increase in the computational costs as the compression becomes more strict.

**Table 3.** One-sided t-test results between different values of hidden layers and shape in the grid search space

| Sample Group | Comparison Group | Bottleneck = 2 | | Bottleneck = 3 | |
|---|---|---|---|---|---|
| Number of Hidden Layers | | t-Statistic | $p$-value | t-Statistic | $p$-value |
| 5 | 4 | -1.34 | 0.91 | -1.31 | 0.90 |
| | 3 | -1.00 | 0.84 | -1.22 | 0.89 |
| | 2 | 0.13 | 0.45 | -0.79 | 0.78 |
| | 1 | 3.48 | $3.09 \times 10^{-4}$ (***) | 0.40 | 0.34 |
| 4 | 3 | 4.07 | $3.34 \times 10^{-5}$ (***) | 2.28 | 0.01 (*) |
| | 2 | 10.96 | $1.43 \times 10^{-22}$ (***) | 11.32 | $4.08 \times 10^{-22}$ (***) |
| | 1 | 19.60 | $5.39 \times 10^{-49}$ (***) | 20.29 | $5.08 \times 10^{-45}$ (***) |
| 3 | 2 | 9.81 | $3.79 \times 10^{-19}$ (***) | 11.63 | $6.09 \times 10^{-23}$ (***) |
| | 1 | 19.13 | $1.28 \times 10^{-47}$ (***) | 21.41 | $1.26 \times 10^{-47}$ (***) |
| 2 | 1 | 18.11 | $1.50 \times 10^{-44}$ (***) | 16.30 | $3.16 \times 10^{-35}$ (***) |
| Network Shape | | | | | |
| rectangular | elliptic | -0.60 | 0.73 | -1.09 | 0.14 |

One-tailed t-tests were conducted using the grid search results of all the haplotype blocks which had 5 hidden layers in their optimal settings, separately for the bottleneck sizes of 2 and 3. For each block, keeping the shape fixed, we calculated the difference in the validation SNP accuracy when the number of hidden layers increases, for instance from 4 to 5. Then, we ran a one-tailed t-test to examine whether the residuals are significantly greater than zero. We repeated this procedure for each pair of different hidden layer values, as well as between two network shapes to test the statistical advantage of the rectangular autoencoders over the elliptic ones.

(*): $p - value \leq 0.05$

(***): $p - value \leq 0.001$

**Table 4.** Results of the Autoencoder Compression and Reconstruction of Chromosome 22

| Performance Metric | Train | | Test | |
|---|---|---|---|---|
| | Mean | Std. Dev. | Mean | Std. Dev. |
| MSE Loss | $5.69 \times 10^{-3}$ | $1.94 \times 10^{-2}$ | $4.75 \times 10^{-3}$ | $7.84 \times 10^{-3}$ |
| SNP Accuracy | 99.55% | $7.24 \times 10^{-3}$ | 99.56% | $7.08 \times 10^{-3}$ |
| SNP Accuracy for 0s | 99.95% | $1.60 \times 10^{-3}$ | 99.95% | $1.59 \times 10^{-3}$ |
| SNP Accuracy for 1s | 98.59% | $2.56 \times 10^{-2}$ | 98.63% | $2.52 \times 10^{-2}$ |
| SNP Accuracy for 2s | 99.48% | $1.83 \times 10^{-2}$ | 99.50% | $1.83 \times 10^{-2}$ |
| SNP Accuracy for Cases | 99.54% | $7.22 \times 10^{-3}$ | 99.57% | $7.01 \times 10^{-3}$ |
| SNP Accuracy for Controls | 99.55% | $7.25 \times 10^{-3}$ | 99.56% | $7.10 \times 10^{-3}$ |

The means and standard deviations of MSE losses and SNP reconstruction accuracies were computed for 2854 autoencoders utilized for compression of all the haplotype blocks in Chromosome 22. The statistics of SNP accuracies were also reported separately for 3 categories of genotype dosage values in the data (0s, 1s and 2s), and for 2 phenotypic groups (ALS patients and healthy controls).

**Table 5.** Results of the PCA Compression and Reconstruction of Chromosome 22

| Performance Metric | Train | | Test | |
|---|---|---|---|---|
| | Mean | Std. Dev. | Mean | Std. Dev. |
| MSE Loss | $1.54 \times 10^{-2}$ | $1.53 \times 10^{-2}$ | $1.48 \times 10^{-2}$ | $1.50 \times 10^{-2}$ |
| SNP Accuracy | 98.32% | $1.79 \times 10^{-2}$ | 98.39% | $1.75 \times 10^{-2}$ |
| SNP Accuracy for 0s | 99.10% | $1.18 \times 10^{-2}$ | 99.13% | $1.15 \times 10^{-2}$ |
| SNP Accuracy for 1s | 96.42% | $4.50 \times 10^{-2}$ | 96.55% | $4.42 \times 10^{-2}$ |
| SNP Accuracy for 2s | 96.51% | $5.47 \times 10^{-2}$ | 96.69% | $5.35 \times 10^{-2}$ |
| SNP Accuracy for Cases | 98.31% | $1.79 \times 10^{-2}$ | 98.42% | $1.74 \times 10^{-2}$ |
| SNP Accuracy for Controls | 98.32% | $1.79 \times 10^{-2}$ | 98.39% | $1.75 \times 10^{-2}$ |

The means and standard deviations of MSE losses and SNP reconstruction accuracies were computed for PCA transformation and reconstruction of all the 2854 haplotype blocks in Chromosome 22. The number of principal components computed for each block is equal to the bottleneck width designated for its associated autoencoder. The statistics of SNP accuracies were also reported separately for 3 categories of genotype dosage values in the data (0s, 1s and 2s), and for 2 phenotypic groups (ALS patients and healthy controls).

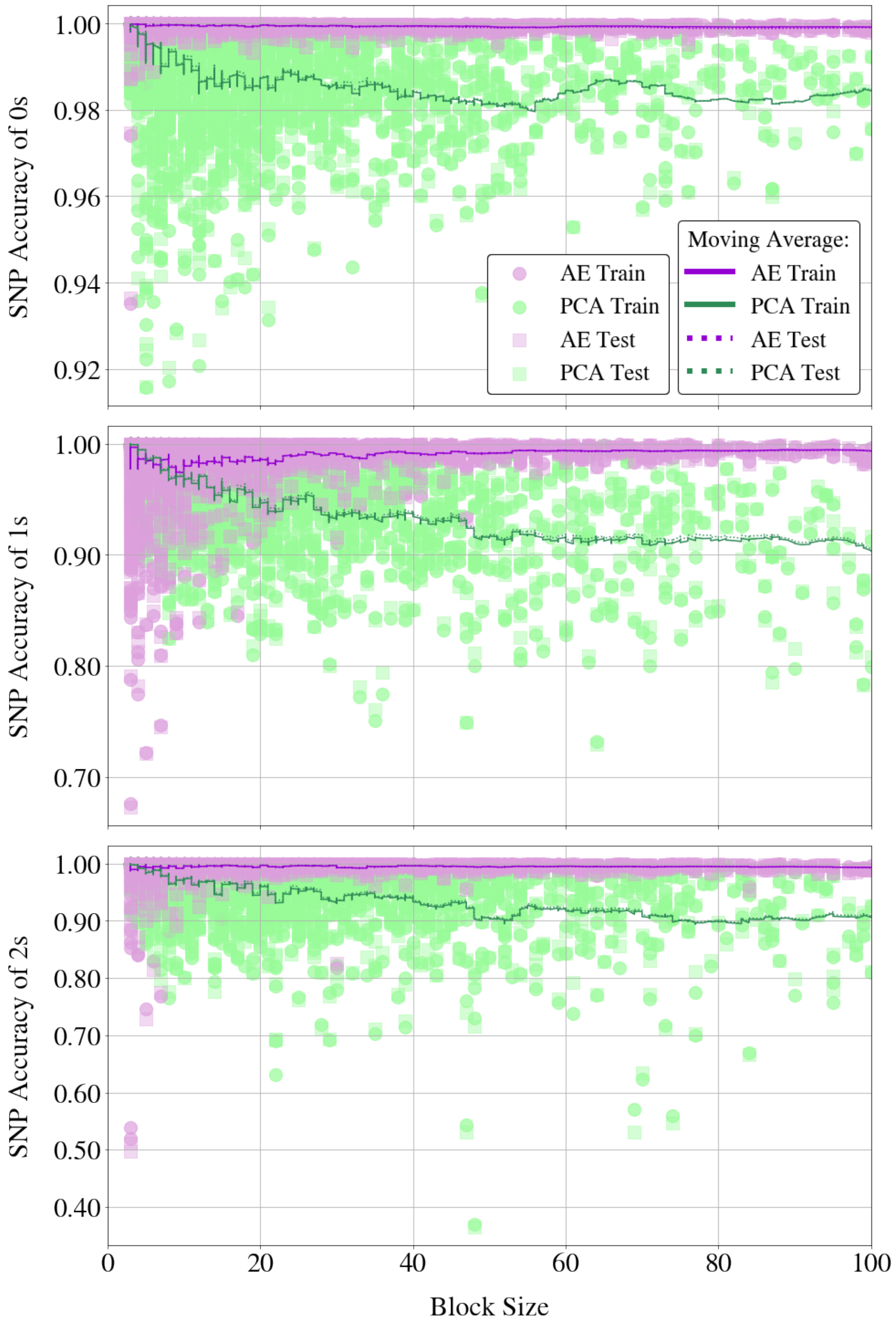


Fig. 11: Scatter plots of SNP reconstruction accuracies obtained by both AE and PCA for allelic dosage values of 0, 1, and 2, from top to bottom. The $x$-axes represent block sizes up to 100 SNPs, shared across all plots. Solid lines illustrate the moving average of SNP accuracies for 2 methods on training samples using a window size of 50, while dotted lines depict the same for test accuracies.

Supplementary note 1

**Project MinE ALS GWAS Consortium authors**

Wouter van Rheenen[1], Mark K. Bakker[1], Kristel R. van Eijk[1], Maarten Kooyman[1], Ahmad Al Khleifat[2], Alfredo Iacoangeli[2,60,81], Nicola Ticozzi[3,61], Johnathan Cooper-Knock[4], Marta Gromicho[5], Siddharthan Chandran[6,62], Karen E. Morrison[7], Pamela J. Shaw[4], John Hardy[8], Michael Sendtner[9], Thomas Meyer[10], Nazli Başak[11], Isabella Fogh[2], Adriano Chiò[12,63], Andrea Calvo[12,63], Elisabetta Pupillo[13], Giancarlo Logroscino[14], Marc Gotkine[15,64], Patrick Vourc'h[16,65], Philippe Corcia[17,65], Philippe Couratier[18,66], Stéphanie Millecamps[19], François Salachas[20,19], Jesus S. Mora Pardina[21], Ricardo Rojas-García[22], Patrick Dion[23], Jay P. Ross[23,67], Albert C. Ludolph[24], Jochen H. Weishaupt[25], Axel Freischmidt[24,68], Gilbert Bensimon[26,69,82,89], Lukas Tittmann[27], Wolfgang Lieb[27], Andre Franke[28], Stephan Ripke[29,70,83], David C. Whiteman[30], Catherine M. Olsen[30], Andre G. Uitterlinden[31,32], Albert Hofman[32], Philippe Amouyel[33], Bryan Traynor[34,71], Adrew B. Singleton[35], Miguel Mitne Neto[36], Ruben J. Cauchi[37], Roel A. Ophoff[38,72,84], Vivianna M. van Deerlin[39], Julian Grosskreutz[40,73], Caroline Graff[41], Lev Brylev[42,74,85], Boris Rogelj[43,75,86], Blaž Koritnik[44], Janez Zidar[44], Zorica Stević[45], Vivian Drory[46,76], Monica Povedano[47], Ian P. Blair[48], Matthew C. Kiernan[49], Garth A. Nicholson[48,77,87], Anjali K. Henders[50], Mamede de Carvalho[5], Susana Pinto[5], Susanne Petri[51], Markus Weber[52], Guy A. Rouleau[23], Vincenzo Silani[3,61], Jonathan Glass[53], Robert H. Brown[54], John E. Landers[54], Christopher E. Shaw[2], Peter M. Andersen[55], Fleur C. Garton[50], Allan F. McRae[50], Russell L. McLaughlin[56], Orla Hardiman[57], Kevin P. Kenna[58,1], Naomi R. Wray[50,78], Ammar Al-Chalabi[2,79], Philip Van Damme[59,80,88], Leonard H. van den Berg[1], Jan H. Veldink[1].

1. Department of Neurology, UMC Utrecht Brain Center, University Medical Center Utrecht, Utrecht University, Utrecht, The Netherlands.
2. Maurice Wohl Clinical Neuroscience Institute, Department of Basic and Clinical Neuroscience, Institute of Psychiatry, Psychology Neuroscience, King's College London, London, UK.
3. Department of Neurology-Stroke Unit and Laboratory of Neuroscience, Istituto Auxologico Italiano IRCCS, Milan, Italy.
4. Sheffield Institute for Translational Neuroscience (SITraN), University of Sheffield, Sheffield, UK.
5. Instituto de Fisiologia, Instituto de Medicina Molecular João Lobo Antunes, Faculdade de Medicina, Universidade de Lisboa, Lisbon, Portugal.
6. Euan MacDonald Centre for Motor Neurone Disease Research, Edinburgh, UK.
7. School of Medicine, Dentistry, and Biomedical Sciences, Queen's University Belfast, Belfast, UK.
8. Department of Molecular Neuroscience, Institute of Neurology, University College London, London, UK.
9. Institute of Clinical Neurobiology, University Hospital Würzburg, Würzburg, Germany.
10. Charité University Hospital, Humboldt-University, Berlin, Germany.
11. Neurodegeneration Research Laboratory, Boğaziçi University, Istanbul, Turkey.
12. "Rita Levi Montalcini" Department of Neuroscience, ALS Centre, University of Torino, Turin, Italy.
13. Research Center for ALS, Department of Neuroscience, Laboratory of Neurological Diseases, Istituto di Ricerche Farmacologiche Mario Negri IRCCS, Milan, Italy.
14. Department of Clinical Research in Neurology, University of Bari at "Pia Fondazione Card G. Panico" Hospital, Bari, Italy.
15. Faculty of Medicine, Hebrew University of Jerusalem, Israel.
16. Service de Biochimie et Biologie moléculaire, CHU de Tours, Tours, France.
17. Centre de référence sur la SLA, CHU de Tours, Tours, France.
18. Centre de référence sur la SLA, CHRU de Limoges, Limoges, France.
19. ICM, Institut du Cerveau, Inserm, CNRS, Sorbonne Université, Hôpital Pitié-Salpêtrière, Paris, France.
20. Département de Neurologie, Centre de référence SLA Ile de France, Hôpital de la Pitié Salpêtrière, AP-HP, Paris, France.
21. ALS Unit, Hospital San Rafael, Madrid, Spain.
22. MND Clinic, Neurology Department, Hospital de la Santa Creu i Sant Pau de Barcelona, Universitat Autonoma de Barcelona, Barcelona, Spain.
23. Montreal Neurological Institute and Hospital, McGill University, Montréal H3A 2B4, Canada.
24. Department of Neurology, Ulm University, Ulm, Germany.
25. Division of Neurodegeneration, Department of Neurology, University Medicine Mannheim, Medical Faculty Mannheim, Heidelberg University, Mannheim, Germany.
26. Département de Pharmacologie Clinique, Hôpital de la Pitié-Salpêtrière, UPMC Pharmacologie, AP-HP, Paris, France.
27. Popgen Biobank and Institute of Epidemiology, Christian Albrechts-University Kiel, Kiel, Germany.
28. Institute of Clinical Molecular Biology, Kiel University, Kiel, Germany.
29. Analytic and Translational Genetics Unit, Massachusetts General Hospital, Boston, Massachusetts, USA.
30. Cancer Control Group, QIMR Berghofer Medical Research Institute, Herston, QLD, Australia.
31. Department of Internal Medicine, Genetics Laboratory, Erasmus Medical Center Rotterdam, Rotterdam, The Netherlands.
32. Department of Epidemiology, Erasmus Medical Center Rotterdam, Rotterdam, The Netherlands.
33. Univ. Lille, Inserm, Centre Hosp. Univ. Lille, Institut Pasteur de Lille, UMR1167 - RID-AGE LabEx DISTALZ - Risk factors and molecular determinants of aging-related diseases, F-59000 Lille, France.
34. Neuromuscular Diseases Research Section, Laboratory of Neurogenetics, National Institute on Aging, NIH, Porter Neuroscience Research Center, Bethesda, Maryland, USA.
35. Molecular Genetics Section, Laboratory of Neurogenetics, National Institute on Aging, NIH, Porter Neuroscience Research Center, Bethesda, Maryland, USA.

36. Universidade de São Paulo, São Paulo, Brazil.
37. Centre for Molecular Medicine and Biobanking Department of Physiology and Biochemistry, Faculty of Medicine and Surgery, University of Malta, Malta.
38. University Medical Center Utrecht, Department of Psychiatry, Rudolf Magnus Institute of Neuroscience, The Netherlands
39. Center for Neurodegenerative Disease Research, Perelman School of Medicine at the University of Pennsylvania, Philadelphia, Pennsylvania, USA.
40. Hans-Berger-Department of Neurology, Jena University Hospital, Jena, Germany.
41. Department of Geriatric Medicine, Karolinska University Hospital-Huddinge, Stockholm, Sweden.
42. Department of Neurology, Bujanov Moscow Clinical Hospital, Moscow, Russia.
43. Department of Biotechnology, Jožef Stefan Institute, Ljubljana, Slovenia.
44. Ljubljana ALS Centre, Institute of Clinical Neurophysiology, University Medical Centre Ljubljana, Ljubljana, Slovenia.
45. Clinic of Neurology, Clinical Center of Serbia, School of Medicine, University of Belgrade, Belgrade, Serbia.
46. Neuromuscular Diseases Unit, Department of Neurology, Tel Aviv Sourasky Medical Center, Tel Aviv, Israel.
47. Functional Unit of Amyotrophic Lateral Sclerosis (UFELA), Service of Neurology, Bellvitge University Hospital, L'Hospitalet de Llobregat, Barcelona, Spain.
48. Centre for Motor Neuron Disease Research, Faculty of Medicine, Health and Human Sciences, Macquarie University, NSW 2109, Australia.
49. Brain and Mind Centre, University of Sydney, Sydney, New South Wales, Australia.
50. Institute for Molecular Bioscience, University of Queensland, Brisbane, Queensland, Australia.
51. Department of Neurology, Hannover Medical School, Hannover, Germany.
52. Neuromuscular Diseases Unit/ALS Clinic, Kantonsspital St. Gallen, 9007, St. Gallen, Switzerland.
53. Department Neurology, Emory University School of Medicine, Atlanta, Georgia, USA.
54. Department of Neurology, University of Massachusetts Medical School, Worcester, Massachusetts, USA.
55. Department of Clinical Sciences, Neurosciences, Umeå University, SE-901 85 Umeå, Sweden.
56. Complex Trait Genomics Laboratory, Smurfit Institute of Genetics, Trinity College Dublin, Dublin D02 PN40, Ireland.
57. Academic Unit of Neurology, Trinity Biomedical Sciences Institute, Trinity College Dublin, Dublin D02 PN40, Ireland.
58. Department of Translational Neuroscience, UMC Utrecht Brain Center, University Medical Center Utrecht, Utrecht University, Utrecht, The Netherlands.
59. KU Leuven – University of Leuven, Department of Neurosciences, Experimental Neurology, and Leuven Brain Institute (LBI), Leuven, Belgium.
60. Department of Biostatistics and Health Informatics, Institute of Psychiatry, Psychology and Neuroscience, King's College London, London, UK.
61. Department of Pathophysiology and Transplantation, "Dino Ferrari" Center, Università degli Studi di Milano, Milan, Italy.
62. Centre for Neuroregeneration and Medical Research Council Centre for Regenerative Medicine, University of Edinburgh, Edinburgh, UK.
63. Neurologia 1, Azienda Ospedaliero Universitaria Città della Salute e della Scienza, Turin, Italy.
64. The Agnes Ginges Center for Human Neurogenetics, Dept. of Neurology, Hadassah Medical Center, Jerusalem, Israel.
65. UMR 1253, Université de Tours, Inserm, 37044 Tours, France.
66. UMR 1094, Université de Limoges, Inserm, 87025 Limoges, France.
67. Department of Human Genetics, McGill University, Montreal, QC H3A 0C7, Canada.
68. German Center for Neurodegenerative Diseases (DZNE) Ulm, Ulm, Germany.
69. Pharmacologie Sorbonne Université, Paris, France.
70. Stanley Center for Psychiatric Research, Broad Institute of MIT and Harvard, Cambridge, Massachusetts, USA.
71. Department of Neurology, Johns Hopkins University, Baltimore, Maryland, USA.
72. Department of Human Genetics, David Geffen School of Medicine, University of California, Los Angeles, California, USA.
73. Precision Neurology Unit, Department of Neurology, University Hospital Schleswig-Holstein, University of Luebeck, Luebeck, Germany.
74. Moscow Research and Clinical Center for Neuropsychiatry of the Healthcare Department, Moscow, Russia.
75. Biomedical Research Institute BRIS, Ljubljana, Slovenia.
76. Faculty of Medicine, Tel Aviv University, Tel Aviv, Israel.
77. Northcott Neuroscience Laboratory, ANZAC Research Institute, Concord, NSW 2139, Australia.
78. Queensland Brain Institute, University of Queensland, Brisbane, Queensland, Australia.
79. King's College Hospital, Denmark Hill, SE5 9RS London, UK.
80. VIB, Center for Brain Disease Research, Laboratory of Neurobiology, Leuven, Belgium.
81. National Institute for Health Research Biomedical Research Centre and Dementia Unit, South London and Maudsley NHS Foundation Trust and King's College London, London, UK.
82. Institut du Cerveau, Paris Brain Institute ICM, Paris, France.
83. Department of Psychiatry and Psychotherapy, Charité - Universitätsmedizin, Berlin 10117, Germany.
84. Center for Neurobehavioral Genetics, Semel Institute for Neuroscience and Human Behavior, University of California, Los Angeles, California, USA.

85. Department of Functional Biochemistry of the Nervous System, Institute of Higher Nervous Activity and Neurophysiology Russian Academy of Sciences, Moscow, Russia.
86. Faculty of Chemistry and Chemical Technology, Universtiy of Ljubljana, Ljubljana, Slovenia.
87. Molecular Medicine Laboratory, Concord Repatriation General Hospital, Concord, NSW 2139, Australia.
88. University Hospitals Leuven, Department of Neurology, Leuven, Belgium.
89. Laboratoire de Biostatistique, Epidémiologie clinique, Santé Publique Innovation et Méthodologie (BESPIM), CHU-Nîmes, Nîmes, France.
